# Attention enhances LFP phase coherence in macaque visual cortex, improving sensory processing

**DOI:** 10.1101/499756

**Authors:** Behzad Zareian, Mohammad Reza Daliri, Kourosh Maboudi, Hamid Abrishami Moghaddam, Stefan Treue, Moein Esghaei

## Abstract

Attention selectively routes the most behaviorally relevant information among the vast pool of sensory inputs through cortical regions. Previous studies have shown that visual attention samples the surrounding stimuli periodically. However, the neural mechanism underlying this sampling in the sensory cortex, and whether the brain actively uses these rhythms, has remained elusive. Here, we hypothesize that selective attention controls the phase of oscillatory synaptic activities to efficiently process the relevant information in the brain. We document an attentional modulation of pre-stimulus inter-trial phase coherence (a measure of deviation between instantaneous phases of trials) at low frequencies in macaque visual area MT. Our data reveal that phase coherence increases when attention is deployed towards the receptive field of the recorded neural population. We further show that the attentional enhancement of phase coherence is positively correlated with the attentional modulation of stimulus induced firing rate, and importantly, a higher phase coherence leads to a faster behavioral response. Our results suggest a functional utilization of intrinsic neural oscillatory activities for better processing upcoming environmental stimuli, generating the optimal behavior.

## Introduction

One of the most important cognitive functions of the mammalian brain is selective attention. Attention selectively routes the most behaviorally relevant information among the vast pool of sensory inputs through cortical regions. This allows the brain to compensate the limited neural resources and create appropriate behavioral responses quickly (Petersen and Posner, 2012).

Attentional influences on neural responses in sensory regions have been extensively documented; effects which reflect a multitude of aspects of cortical information processing (Baluch and Itti, 2011; Maunsell and Treue, 2006; Petersen and Posner, 2012; Reynolds and Chelazzi, 2004). Covertly directing attention towards the receptive field of a visual neuron enhances the neural responses even in the absence of visual stimulation (Kastner et al., 1999; Treue, 2001), alters the shape and form of receptive fields (Niebergall et al., 2011b, 2011a; Womelsdorf et al., 2008), modulates the variability and temporal structure of the neuron’s firing patterns (Mitchell et al., 2009; Xue et al., 2017), modulates inter-neuronal correlations to increase neural discriminability (Cohen and Maunsell, 2009; Esghaei et al., 2015a) and synchronizes neighboring neurons, presumably to better propagate information to downstream areas (Fries et al., 2008; Siegel et al., 2008).

Attention has been suggested to exploit oscillatory neural activities, as of oscillatory components of local field potentials (LFP), to enhance the efficacy of cortical processing (Fries, 2009; Harris and Thiele, 2011; Khayat et al., 2010; Lakatos et al., 2008; Noudoost et al., 2010). LFPs represent synaptic activities of local cortical neuronal populations (Einevoll et al., 2013). Their oscillations are tightly linked to attention in both low and high frequencies (Chalk et al., 2010; Fan et al., 2007; Fries et al., 2001; Khayat et al., 2010; Mo et al., 2011). For instance, Fries et al. showed that synchronization in the gamma band increases for neurons activated by the attended stimulus (Fries et al., 2001). Moreover, recent investigations document a prominent role of low frequency oscillations, especially in the alpha band, in attentional processing and shaping large-scale task-related functional networks of the brain (de Pesters et al., 2016; Helfrich et al., 2018; Van Kerkoerle et al., 2014). Alpha oscillations provide periodic alternations in the neural excitability, causing a rhythmic modulation of perception (Haegens et al., 2015; Jensen et al., 2014; Jensen and Mazaheri, 2010; Kizuk and Mathewson, 2017; Lorincz et al., 2009; Milton and Pleydell-Pearce, 2016). Similarly alpha amplitude in human cortex has been shown to be manipulated by attention to modulate neural processing (Yamagishi et al., 2003). Although there is prominent evidence on attentional modulation of low frequency amplitude, the role of low frequency phase in attentional processing is yet controversial.

The phase of low frequency oscillations modulates local neural activities represented by gamma band activity, which presumably enables distant brain regions to interact (Demiralp et al., 2007). Some studies have shown that the phase of ongoing neural oscillations is responsible for periodic sampling by visual attention (Busch and VanRullen, 2010; VanRullen et al., 2011). Furthermore, the phase of low frequency oscillations facilitates information transfer and neural coding in the brain (Voloh and Womelsdorf, 2016). Therefore, low frequency phase can enable the neural system to prepare for processing upcoming sensory stimuli.

Pre-stimulus neural activity has been shown to be a determinant of retrieving episodic memory, perception of environmental information and attention-related variability in response speed (Addante et al., 2011; Hanslmayr et al., 2013, 2007; Shibata et al., 2008). Interestingly, it has been shown that pre-stimulus brain activity causally determines visual perception (Dugué et al., 2011). In addition, it has been shown that the phase of low frequency oscillations is responsible for this causal relationship (Hanslmayr et al., 2013). Furthermore, attention has been reported to determine the phase of low frequency neural oscillations in order to influence neuronal responses and behavioral responses to external events (Lakatos et al., 2008). Lakatos et al. (2008) showed that V1 neurons are phase-locked to those rhythmically presented stimuli that are attended, presumably to generate a larger evoked response and induce faster behavioral response times (Lakatos et al., 2008). Nevertheless, it is not clear whether spontaneous, rather than externally induced low frequency neural activities are harnessed by attention. Given the prominent role of low frequency phase in shaping perception, we hypothesize that selective attention should control the LFP phase, potentially to route information in the brain.

Here, we investigate the influence of attention on the phase of pre-stimulus low frequency oscillations. We calculate the effect of attention on inter-trial phase coherence (a measure of deviation between instantaneous phases of trials) at low frequencies in area MT. Our results reveal that phase coherence increases when attention is deployed towards the RF of the recorded neuron. We further show that higher phase coherence leads to shorter reaction times and phase coherence modulation correlates positively with attentional modulation of firing rate after stimulus onset, together suggesting that phase coherence is a tool that attention exploits to generate an optimal visuo-motor response.

## Results

Two monkeys were trained and cued to covertly direct their spatial attention to one of two moving random dot patterns (RDP), each presented in one hemifield. Within the cued pattern, they had to detect a brief change in the color or motion direction. To initiate a trial, the monkeys touched a lever and foveated a central fixation spot presented on the screen. After 150 ms, the cue, either a small static colored RDP or a moving RDP, appeared on either side (near the fixation spot), indicating the location of the upcoming target stimulus. The cue disappeared after 500 ms, followed by two moving RDPs, one placed inside the receptive field of the neuron being recorded and the other on the opposite side of the fixation point (Figure 1 A). The animals had to report a brief color/direction change in the target, ignoring changes in the distracter. We recorded single unit and LFP signals from 11 and 29 sites in the visual area MT of each monkey, while they carried out the task. Both animals showed a high behavioral performance. Overall, target detection rates for color and direction tasks were 91% and 88%, respectively. Monkey C correctly ignored the distractor in 85% and monkey T in 91% of the trials. Monkey C & T’s average behavioral reaction time was 392 ms and 351 ms, respectively (see the details of the behavioral paradigm and recording in ‘Experimental Procedures’).

**Figure 1.**
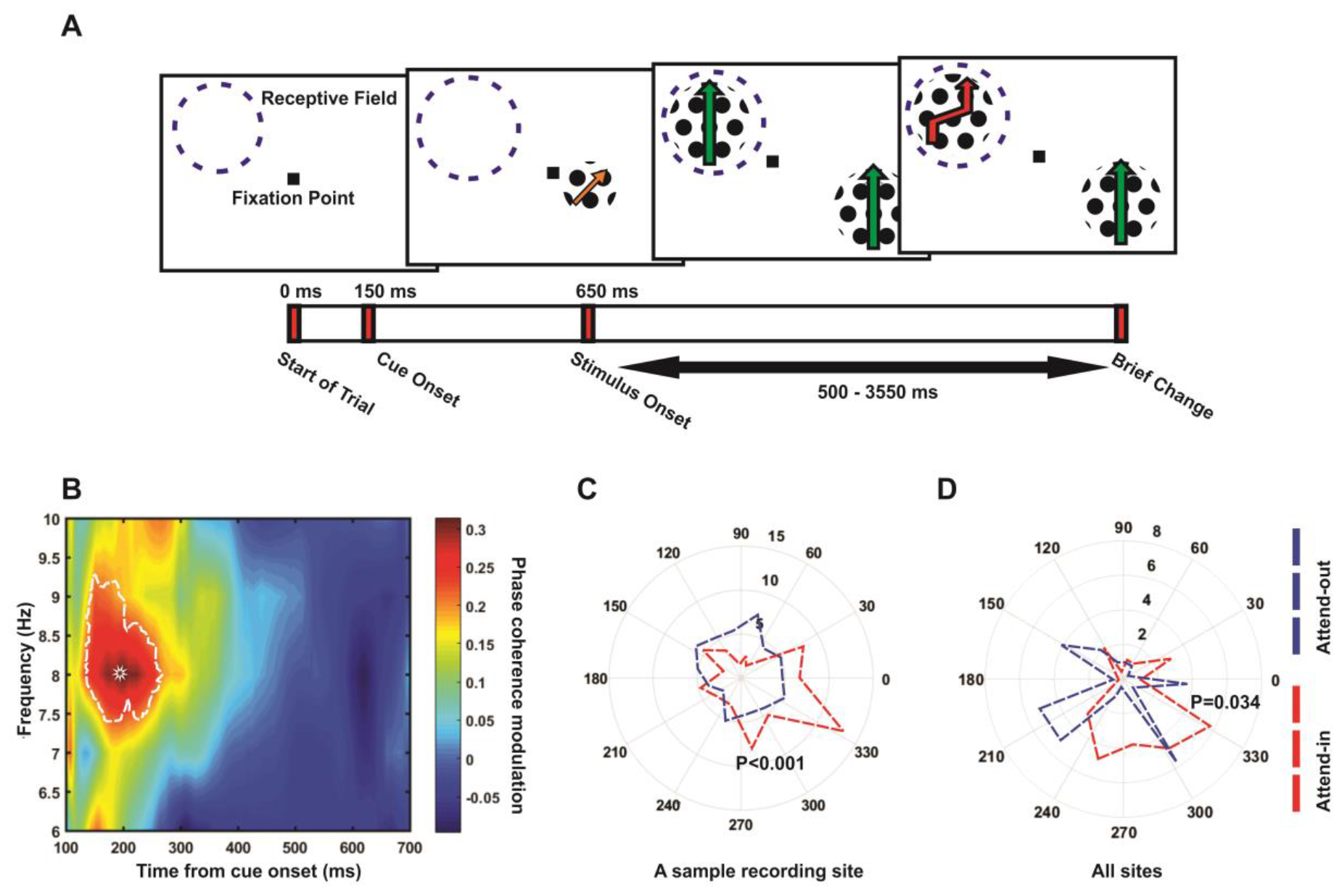
Attention modulates phase coherence. (A) Behavioral paradigm. Each trial started when the monkey foveated a central fixation point and touched a lever. The receptive field of the neuron under study is indicated by a dashed circle (not present on the screen). (B) Phase coherence modulation (PCM) Map. X-axis plots time (ms) aligned to the cue onset and Y-axis represents the LFP oscillation frequency in Hz. Each Y-axis value indicates the center of a 4 Hz frequency band. The colors represent the values of the PCM calculated by the formula: (attend-in phase coherence - attend-out phase coherence) / (attend-in phase coherence + attend-out phase coherence), averaged across the 31 sites with at least 50 trials. The region indicated by the saturated line shows frequency-time pairs with a statistically significant PCM. A star marks the frequency-time pair with the maximum PCM (at 200 ms, 8 Hz) (C) Polar histogram of the instantaneous phase for a sample site in attend-in (red) and attend-out (blue) trials for the time-frequency pair (200 ms, 8 Hz). The values indicate the number of trials that share a given phase. A high value therefore indicates a large amount of inter-trial coherence for that instantaneous LFP phase. The total number of trials are 70 for each condition in this site. As the figure shows, the trials in the attend-in condition are more coherent (towards phase 330°) than trials in the attend-out condition (p-value <0.001 for attend-in condition, p-value=0.9 for attend-out condition; Rayleigh test, p-value<0.001 for difference in phase coherence; permutation test) (D) Histograms of average phases of recording sites separated by the attention condition. The polar histograms consist of 15 bins and in each bin, the number of average phase vectors of sub-trials (separated by attention condition) in sites is plotted (p-value <0.0001 for attend in sites’ phases, p-value=0.12 for attend out ones; Rayleigh test. p-value=0.034 for difference in the coherence of the sites’ average phases between attention conditions; permutation test). The mean vectors of the attend-in and attend-out groups are shown in red and blue, respectively (See also Figure S1, Figure S2, Figure S3 and Figure S4).

To investigate if attention induces any preparatory neural activity before processing upcoming visual stimuli, we analyzed the Local field potentials (LFPs) following 100 ms after cue onset and before the onset of the RDPs, a time window without a stimulus in the receptive field (RF). LFPs were filtered in 4 Hz frequency bands stepped by 1 Hz with the middle frequency of the bands changing between 6-10 Hz. For each frequency band, Hilbert transform was used to compute the instantaneous phases of the signals. To investigate if attention modulates the variation of these phases across trials, we computed the inter-trial phase coherence separately for trials in which the monkeys attended to the stimulus inside the receptive field (attend-in condition) and for those trials in which the monkeys attended to the stimulus outside receptive field (attend-out condition). To compute this phase coherence for each attention condition in a given frequency band and time point, the following analysis steps were taken: 1. We imposed unit-length vectors with their phases coming from the trials with the corresponding attention condition at the given frequency band and time. 2. Phase coherence was quantified by calculating the length of the average vector. 3. We calculated the attentional modulation of phase coherence for each frequency band at each time per site using the following formula: (attend-in phase coherence - attend-out phase coherence) / (attend-in phase coherence + attend-out phase coherence). 4. Phase coherence modulations (PCM) were averaged across sites. Figure 1B maps the PCM values starting from 100 ms after the onset of the cue until 700 ms later, in 4 Hz frequency bands with their middle frequencies stepped by 1 Hz from 6 to 10 Hz (4-8 Hz, 5 to 9 Hz, etc.). The colors represent the PCM magnitudes. Those time-frequency pairs with a significant PCM are indicated with a white border (p<0.01; ttest) in the map (See Experimental Procedures for more details). A star marks the position (200 ms, 6-10 Hz) of the highest PCM (31%) across all time points and frequencies (PCM for other frequency ranges as well as the phase coherence for the attend-in and attend-out conditions, separately are shown in Figure S1). As shown in the Figure 1B, there is a cluster of time points across neighboring frequencies centered at 8 Hz, in which attention has enhanced the phase coherence significantly for the attend-in relative to the attend-out condition. A sample site’s phases are presented in Figure 1C, showing that the LFP phase is more densely concentrated in the attend-in subset of trials (red), compared to the attend-out trials (200 ms, 6-10 Hz) (p<0.001; permutation test). Figure 1D shows the distribution of the average phase (over trials) for all sites. The average phases of sites are clearly more coherent in the attend-in than attend out trials at (200 ms, 6-10 Hz) (p=0.034; permutation test). This further indicates that the phase coherence between sites is also enhanced in attend-in, compared to the attend-out condition.

To control for the sensory influence of the cue (assuming that LFP has a larger amplitude in attend-out compared to the attend-in condition, when cue evokes a sensory influence), we separated the sites where cue (shown in attend-in condition) enhanced the LFP amplitude or reduced it (compared to the attend-out condition). This division was made based on the average LFP amplitude in the 100 ms time interval surrounding the time point with the maximum phase coherence modulation (“evoked response control window” of 150-250 ms from cue onset, Figure 2A). Figure 2B shows the average LFP amplitude within the evoked response control window for the two groups of sites. The “with putative sensory evoked response” and “without putative sensory evoked response” groups are depicted in orange and green, respectively. Figure 2C and D show the time-resolved average LFP responses for these two groups, separately. We assume that if the phase coherence modulation effect is simply a side effect of the cue’s evoked sensory response, then the phase coherence modulation should be observed only in sites with a putative sensory evoked response. However, both groups of sites showed a significant phase coherence modulation (“with putative sensory evoked response”: p=0.001- Figure 2E, “without putative sensory evoked response”: p=0.027- Figure 2F; ttest). These results suggest that the phase coherence modulation observed here, is caused by attention, rather than being a side effect of cue’s sensory response.

**Figure 2.**
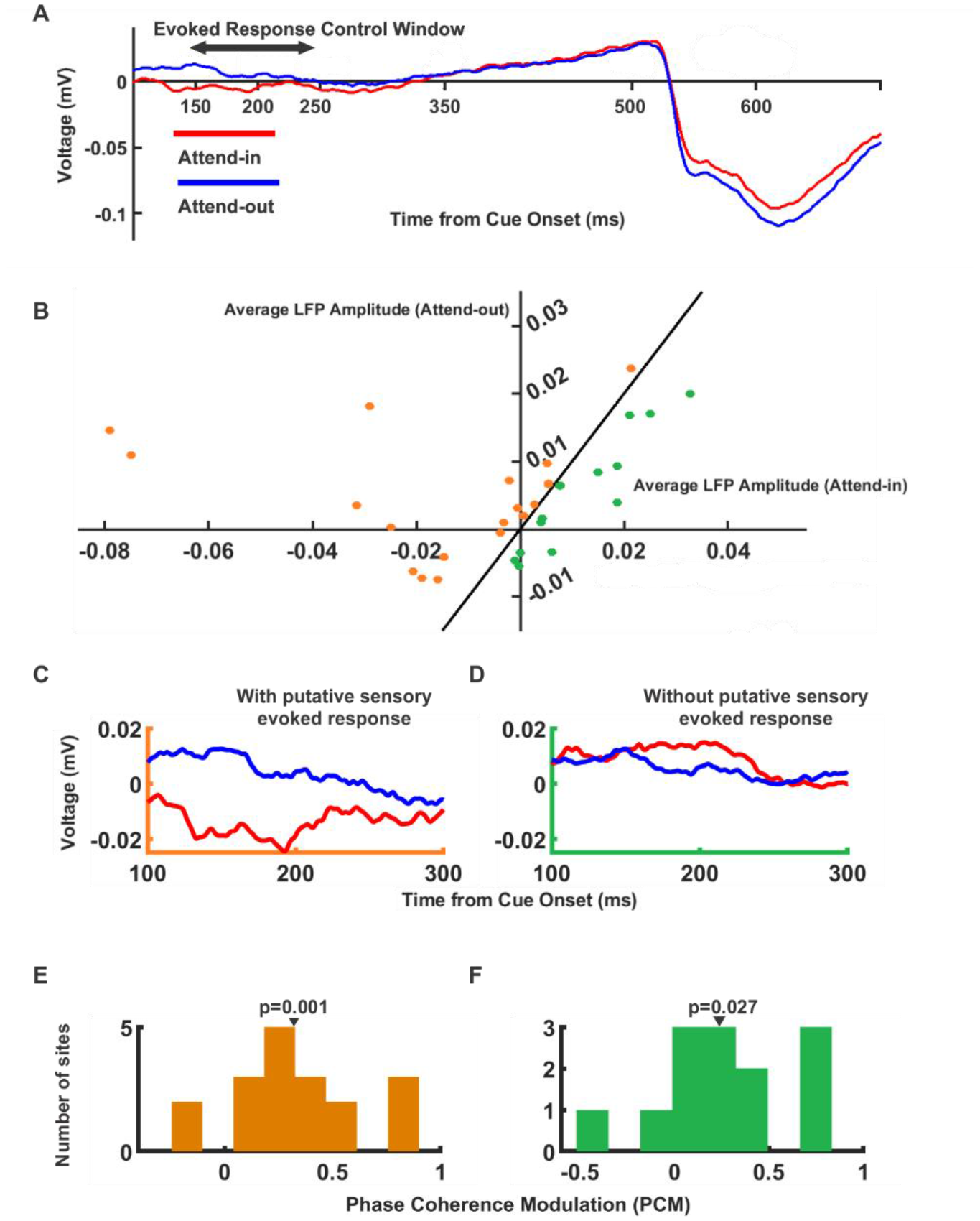
Control for sensory influence of the evoked response on phase coherence calculation. (A) The grand average of LFPs in attend in (red) and attend out (blue) conditions across all sites. The double-headed arrow shows the time interval that is chosen for the evoked response control computations (150 to 250 ms after cue onset). (B) Average LFP amplitude of attend-in subset of trials for each site versus attend-out subset within the evoked response control window. Orange dots indicate the sites that their average LFP over attend-in trials is lower than attend-out (with putative sensory evoked response) and green dots are the remaining sites (without putative sensory evoked response). (C) The time-resolved average LFP amplitude for sites with putative sensory evoked response and (D) without putative sensory evoked response. (E, F) Histograms of phase coherence modulation for the sites with and without sensory evoked response, within the time-frequency point with maximum PCM (200 ms, 8 Hz) (p-value =0.001 for with putative sensory evoked group, p-value=0.027 for without putative sensory evoked group; ttest).

It could be argued that the comparison of phase coherence measurement across attention conditions may be confounded by the differences of signal to noise ratios in the LFPs across conditions. To test this, we separately calculated the power of LFP oscillations within the 6-10 Hz frequency range and compared it between the attention conditions. The attend-in and attend-out conditions showed no significant difference between their spectral power (Figure S2), suggesting that the observed phase coherence modulation is not a side effect of different spectral powers. In addition, it could be possible that phase coherence modulation is a result of a difference in the arousal level between the attend-in and attend-out trials, rather than the location where spatial attention is directed towards. To test this, we analyzed the reaction times (as a quantification of the average arousal level in a trial) in each of the two conditions. No systematic difference was observed between the reaction times of the two conditions (p-value=0.0548 Wilcoxon signed Rank test between sessions’ average reaction times; Attend-in average RT= 357 ms and Attend-out average RT= 354 ms; Figure S3A). We further excluded those sessions where the attend-in condition was faster than the attend-out condition (corresponding to sessions with a higher arousal in the attend-in condition) and recalculated the phase coherence modulation (at 200 ms, 6-10 Hz). The PCM distribution for those sties recorded in these sessions, was still significantly above zero, meaning that even though the reaction time is not lower for attend-in trials, there still exists a positive PCM, hence the PCM is not a side effect of arousal (Figure S3B).

To further investigate if the PCM observed here, is actually involved in the attentional processing of stimuli coming in the future (and correspondingly reaction time to their change), we next asked if PCM is associated to the well-known neural signature of attention, “spike rate enhancement” (Katzner et al., 2009). We calculated the correlation between PCM at (200 ms, 8 Hz) and attentional modulation of firing rate after stimulus onset across sites. Attentional modulation of firing rate was calculated at each time point, using the attentional index given by the formula: (attend-in spike density function - attend-out spike density function) / (attend-in spike density function + attend-out spike density function). We found that phase coherence modulation was positively correlated with the attentional index following stimulus onset (average Pearson R= 0.076, p-value<0.0001 for sites with positive PCM; ttest – Figure 3A; also average Pearson R=0.026, p-value<0.0001 for all sites), in-line with the account that phase coherence may influence the attentional processing of upcoming stimuli. Each point in Figure 3B illustrates the magnitude of correlation between these two measures across sites. Interestingly, both the dynamics of this correlation and that of the attentional index showed an oscillatory regime within the theta band (the spectral maximum at 5.8 Hz (Figure 3B) and 7.32 Hz (Figure 3C) for the correlation and attentional index, respectively). This indicates that a higher phase coherence modulation leads to an enhanced attentional modulation of firing rate, suggesting that attention may functionally exploit phase coherence to enhance the neural representation of upcoming stimuli.

**Figure 3.**
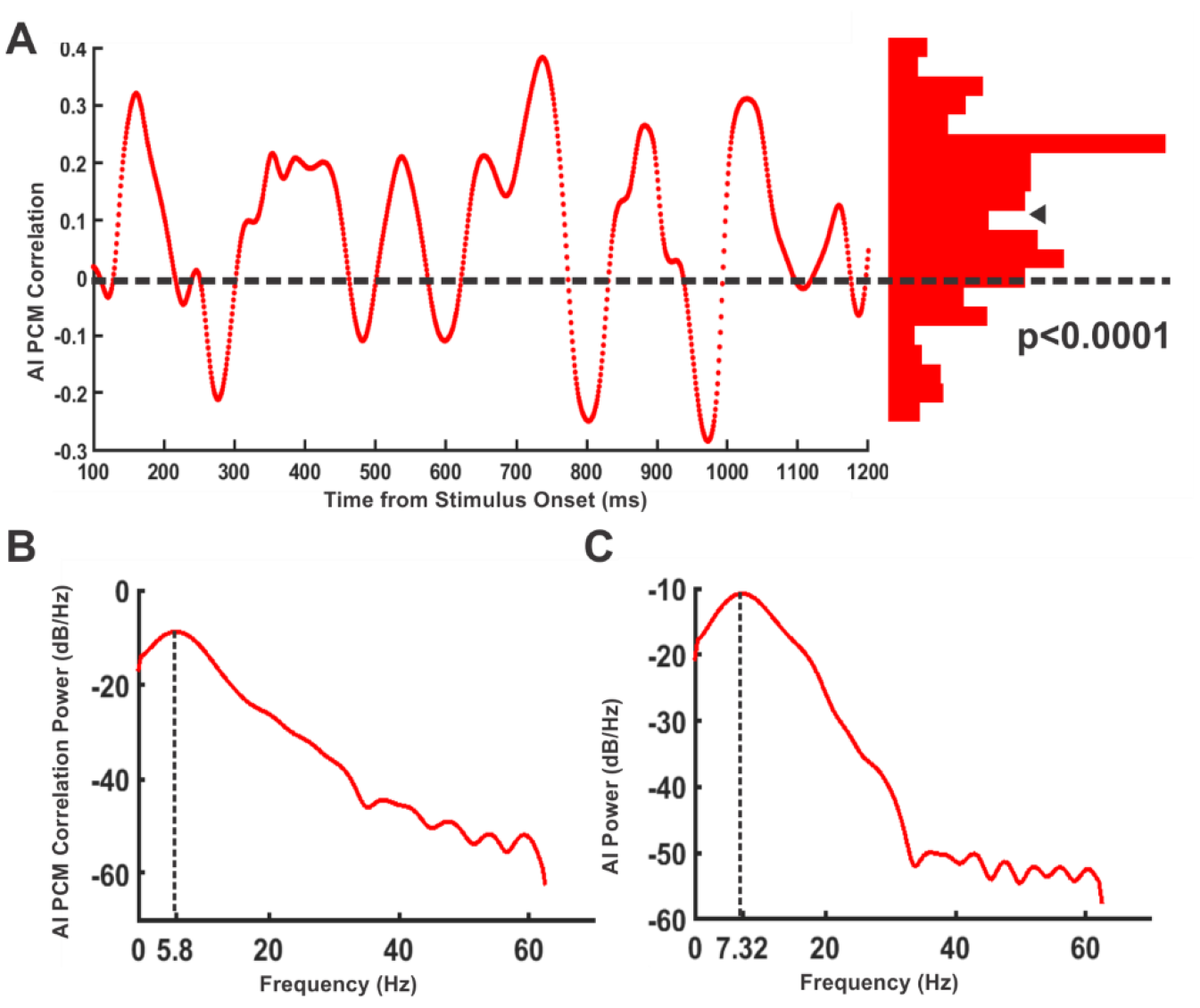
Phase coherence modulation is linked to the neural correlates of attention. (A) Correlation between phase coherence modulation and attentional enhancement of firing rate for time points after stimulus onset (left). The red histogram shows the distribution of correlation magnitudes for different times. The histogram is positively skewed (p-value<0.0001; ttest). (B) Power spectral density of correlation between attentional index and phase coherence modulation (C) Power spectral density of attentional index’s curve of dynamics in time. The frequency with maximum spectral power is indicated by a vertical dashed line.

Our findings suggest that attention resets the phase of low frequency oscillations before the onset of the behaviorally relevant stimulus. This may lead to a more efficient alignment of excitability phases as a preparatory mechanism to better process the upcoming visual stimulus. Therefore, we predict that the monkey’s behavioral performance (as a consequence of neural processing’s efficiency) is linked to phase coherence. We hypothesize that attention shapes sensory processing by modulating inter-trial phase coherence within low frequency oscillatory activities. Given that attention enhances the behavioral detection of stimulus changes (Gonzalez Andino et al., 2005; Prinzmetal et al., 2005), particularly represented by LFPs as long as several seconds before the response event occurs (Parto Dezfouli et al., 2018), we asked if it is the modulation of phase coherence that underlies this attentional influence on behavior.

We conjecture that in those sets of trials with a higher phase coherence, the sensory cortex is prepared more effectively for processing sensory input, leading to a better performance in detecting stimulus changes. We evaluated this by investigating the potential link between phase coherence and the response time of monkeys in reporting the stimulus change. We determined if there is any relationship between how similar a given trial’s phase is to the mean phase, and the reaction time in that trial. The global mean phase (GMP) used in this analysis is the circular average of phases from the trials of all sites with at least 40 trials at the time-frequency pair as the maximum PCM (200 ms, 8 Hz). We expect that in trials where the phase is closer to the GMP, the monkey responds faster. For this step, we analyzed the GMP of attend-in trials due to the higher magnitude of phase coherence among these trials. We observed that the phases of attend-in trials pooled across all sites are significantly biased to the GMP (p-value<0.0001; Rayleigh test). We calculated the correlation between the similarity of phases to GMP and reaction times of the trials and observed that there is a significant negative correlation in the point with the maximum phase coherence modulation (200 ms, 8 Hz) (Pearson’s R=-0.044; p-value<0.001). This shows that the monkey responds faster as the trial’s phase gets closer to the GMP. Further, we grouped the attend-in trials into 16 bins based on their reaction times. We observed a significant negative correlation between the reaction time of the bins and their phase coherence (Pearson’s R=-0.57; p-value=0.02- Figure 4A). To further visualize this, we plotted the phase coherence within the extreme percentiles of the trials according to their response time. Figure 4B illustrates the distribution of vectors for the low and high response time percentiles. The results show that among attend-in trials, the phase coherence of the 5.75 percentile of trials with the smallest response time was higher than that of the 5.75 percentile of trials with the largest response times (p-value<0.0001 for the low response time percentile, p-value=0.42 for the high response time percentile; Rayleigh test, p-value=0.011 for phase coherence difference between the two groups; permutation test). This suggests that phase coherence is a contributing factor to the performance of primates in detecting visual changes, indicating that attention may harness the low frequency phase to improve perception and behavioral responses depending on this perception.

**Figure 4.**
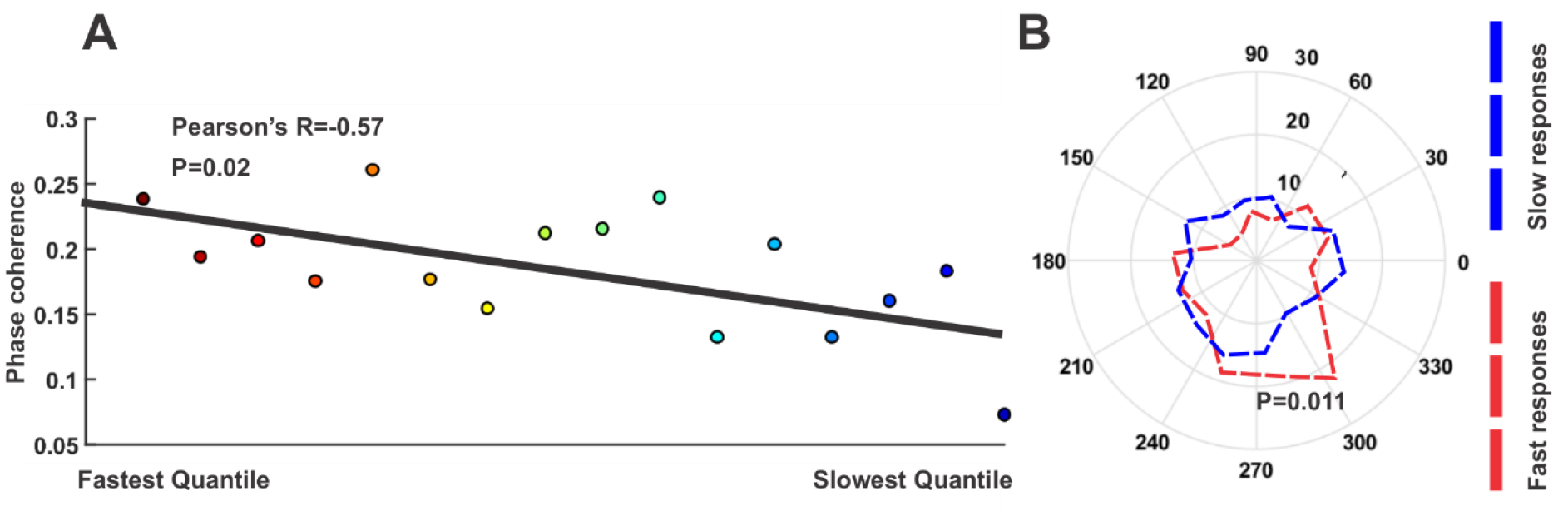
Behavioral correlate of phase coherence modulation. (A) Changes of phase coherence for different subgroups of attend-in trials based on their reaction time. The trials of all sites (31 sites) are divided into 16 distinct subgroups based on their reaction time. There is a negative correlation between the order of these subgroups and their phase coherence (Pearson’s correlation=-0.57, p-value=0.02; ttest) (B) Polar histogram of phases from trials with the longest reaction time vs those with shortest reaction time for the point with the highest PCM (200 ms, 8 Hz). The histogram includes 15 sectors and the number of trial vectors with the longest reaction time is counted in each sector in the most significant point of the PCM map (16th quantile-marked by blue) and the same is counted for the shortest reaction time trials (1st quantile - marked by red). The contours show the distribution of quantiles. Total number of trials in each quantile is 170 (5.75% of all trials in the attend-in condition) (p-value<0.0001 for short response time percentile, p-value=0.42 for long response time percentile; Rayleigh test. p-value=0.011 for difference in phase coherence; permutation test).

## Discussion

It is recently documented that attention is associated with low frequency fluctuations of neural activity. Here, we hypothesized that attention may systematically modulate these low frequency oscillations to enhance the neural representation of upcoming stimuli. To evaluate this, we recorded local field potentials (LFP) from visual area MT of behaving rhesus monkeys, while they performed a visual change detection task, with the focus of their spatial attention directed either inside or outside the receptive field (RF) of the recorded neuron. Our data reveal that switching attention into the RF significantly increases the inter-trial phase coherence within low frequency oscillations around 8 Hz, starting 200 ms after the spatial cue. This suggests for a predictable stimulus, that attention aligns the phase of low frequency oscillatory neural activities to the cue, to optimally prepare processing the stimulus. We further observed that this increase in phase coherence is correlated with the attentional modulation of the single neurons activity, and even the behavioral speed of the animals’ reaction to the stimulus change.

Our main frequency of interest (8 Hz) has been shown to govern endogenous attention (Fiebelkorn et al., 2018, 2013; Helfrich et al., 2018; Landau and Fries, 2012). Importantly, Landau and Fries showed that (1) attention samples multiple stimuli periodically, (2) an attended location is sampled at a frequency of 8 Hz (4 Hz per location) and (3) a cue in one hemifield could reset the attentional sampling temporarily and orient it towards the location of the flash (Landau and Fries, 2012). Here, we document that attention aligns the phase of the oscillatory activity in the same frequency across trials. This suggests that the physiological correlate of the 8 Hz perceptual sampling at the cued location is the phase of LFP at this frequency in a trial. In other words, attention might use the phase of 8 Hz in order to sample the cued location.

But what evidence supports the existence of such a link between the phase of LFP and perceptual sampling? Based on our investigation of reaction times, phase coherence determines subsequent response times of the subject, as a signature of perception efficacy. Similarly, Harris et al. reported that detection of targets depends upon the low frequency phase (in both attended and unattended locations) (Harris et al., 2018). Also, Fiebelkorn et al. documented that the fronto-parietal network’s low frequency fluctuations coordinates the attentional spatial sampling (Fiebelkorn et al., 2018). Furthermore, Busch and VanRullen reported that the visual detection threshold is periodic and strongly correlated with the pre-stimulus phase of EEG signals (Busch and VanRullen, 2010). We further show that attention enhances the perceptual efficacy by aligning the LFP phases preceding the onset of the behaviorally relevant stimulus. This is in line with other recent investigations which relate attention and perception using a similar mechanism (Addante et al., 2011). Furthermore, the steady presentation of stimuli in our attention paradigm, distinguishes it from those studies where the neural activity is entrained to rhythmic stimuli.

Alpha oscillation entrainment has been shown to be maintained and influencing perception even when there is no oscillatory visual stimulation (Spaak et al., 2014). Other studies containing an ongoing oscillatory stimulation, did not clarify if the internally generated (rather than the externally imposed) brain rhythms are influenced by attention. Previous research in the auditory cortex has shown that theta and alpha phase have a profound role in sensory computations, without involving any sensory mediation (Kayser et al., 2016). By using an alternating visual stimulus to induce a neural rhythm, Schroeder and Lakatos showed that attention entrains the phase of the rhythmic neural activity in visual cortex to better process the behaviorally relevant stimulus (Schroeder and Lakatos, 2009). However, their paradigm leaves unclear if attention influences the endogenous oscillatory neural activities. To answer this question, instead of rhythmically presenting the visual stimuli, we presented non-rhythmic stimuli. Our results show that in the absence of any externally evoked neural rhythm, attention modulates the LFP phase preceding the onset of the behaviorally relevant stimulus. Similar to our study, Voloh et al. reported that in the absence of an external sensory entrainment, attentional cues can induce a phase reset which can synchronize high frequency activities and further help the selection of relevant sensory stimuli (Voloh et al., 2015). However, they did not determine whether their observed phase alignment was induced by either the sensory cue or the monkey’s attentive state, rather than selective attention.

We observed the highest attentional modulation of phase coherence within the alpha band. This frequency band has been under investigation in many recent studies, which have shown that alpha band activity inhibits neuronal processing in task-irrelevant areas (Haegens et al., 2011; Klimesch et al., 2007). On the other hand, a decrease in alpha band power can lead to enhanced excitability (Lange et al., 2013). The pre-stimulus alpha phase can change neuronal excitability in order to modify temporal perception, independent from alpha amplitude (Milton and Pleydell-Pearce, 2016). Our study confirms these reports in suggesting that alpha band activity provides a functional tool for selective attention. It can modify the temporal profile of peaks and troughs in the neural activity through phase manipulation, and the spatial profile through changing alpha power. This means that in cortical areas with a larger alpha amplitude, there is more inhibition, and in this way the brain can control a neural population’s potential to suppress activity with attention. Along this line, van Diepen et al. showed that the power and not the phase of alpha oscillations can be modulated by top-down cognitive functions such as attention (van Diepen et al., 2015). However, their study differs from ours in that they examined the phase coherence at the time of target presentation, while we focused on the interval where the monkeys are preparing for the appearance of the behaviorally relevant stimulus. As our results suggest that phase coherence is used as a preparatory mechanism, it is not expected to observe any modulation of it during stimulus presentation. Therefore, our data suggest that the alignment of phase is a tool to prepare the neural system for processing upcoming stimuli, rather than a tool to better process a presented stimulus. Another study has found a similar temporal effect as our finding within the temporal cortex of humans (Yamagishi et al., 2008). They reported that the magnitude of inter-trial coherence increases after cue onset and that phase coherence and performance are positively correlated, consistent with our findings. They speculated that the magnitude of inter-trial coherence could be a measure of attention magnitude among different trials and further suggest that this may reflect neural changes of temporal cortex activity in response to top-down influences. Here we show the first evidence suggesting that attention selectively increases phase coherence in sensory areas that are involved in processing the target stimulus while suppressing it in other cortical areas.

Esghaei et al. showed that switching spatial attention to a neuron’s receptive field reduces the coupling of both gamma oscillations and spikes (both representative of local neural processing) to the low-frequency phase (Esghaei et al., 2015b; Parto Dezfouli et al., 2018) see also (Spyropoulos et al., 2018). These observations may challenge the current finding in that they suggested the coupling of local neural activity to the low frequency phase to have a suppressive role in attention. In the same line, Spyropolous et al. showed that theta rhythms are more prevalent in the attend-out rather than attend in condition (Spyropoulos et al., 2018). However, in the majority of previously used attention paradigms, the stimuli were presented inside the receptive field during the cue period (the period we focused on for phase coherence analyses). Our data on another hand show, in the absence of visual stimulation, that attention exploits the phase of low frequency neural oscillations, potentially to enhance the preparatory mechanisms of visual processing. Correspondingly, when the stimulus appears inside the receptive field and the MT neuron is actively engaged for the task, attention may not use the oscillatory activity anymore. Thus, attention decouples the neurons from the ongoing rhythm to give them the ability to maximize the dissociation of the information contents within the receptive field by making the neurons fire independently (Esghaei et al., 2015b). Meanwhile, it continues to rhythmically sample the other unattended regions by increasing the magnitude of theta at the engaged brain areas. This challenges the notion that attention uses the oscillations always in the same manner. Our results suggest that the function of these oscillations actually depends on the task needs at a given moment; which could be either enhancement of the rhythm for maximizing spatial sampling, or decoupling of neurons from the rhythm’s phase to maximize the neural discrimination within the receptive field.

Inter-areal phase coherence has been proposed to control communication between neighboring brain regions. Zanto et al. showed that alpha-band phase coherence is responsible for longdistance top-down modulation of inter-areal communication by phase-locking separated regions (Zanto et al., 2011). Since an enhancement of inter-trial phase coherence may aid interregional phase coherence (by independently resetting each area’s fluctuations), our finding is in line with the above report, suggesting a neural mechanism by which attention may facilitate the communication of sensory area MT to higher cortical areas. Moreover, it has been shown that the pre-stimulus phase gates information transfer between distant cortical regions (Hanslmayr et al., 2013). The main frequency of their finding (7 Hz) has been shown to synchronize neural outputs among task-related areas which as they point out, can also be described by the communication-through-coherence hypothesis (Fries, 2005; Hanslmayr et al., 2013). This hypothesis suggests that two areas communicate when their excitability states synchronize. On the other hand, low frequency phase can be a tool for coordination of neural oscillations across anatomical and temporal scales during attention. According to Voloh and Womelsdorf, oscillations provide time windows for the optimal transfer of low-level sensory information to higher areas. They suggest that the response to stimuli can be changed dramatically by resetting the oscillations’ periods of excitability to match the presentation of the target stimuli (Voloh and Womelsdorf, 2016). In line with their finding, here we suggest that attention controls the synchronization of MT with higher cortical areas by aligning the low frequency LFP phases of trials for better communication. Nevertheless, further studies need to experimentally examine this using simultaneous recordings from the visual cortex and higher level areas. Also, it may be surprising that the PCM effect observed here, appears only transiently after cue (rather than remaining at an equal magnitude throughout the whole cue period). Considering the fixed and short interval between cue and stimulus onset, the prestimulus period is conceivable to be dominated by the stimulus-locked preparatory activities (as shown in Figure S1). Future investigations may study the stability of this effect by increasing and randomizing the cue interval’s length to remove the preparatory signal.

In summary; we documented a link between attention and phase coherence in low frequencies. Our data show for the first time that attention selectively enhances the inter-trial phase coherence of LFP’s low frequency oscillations in the visual cortex. We further found that higher inter-trial phase coherence leads to an enhanced neural representation and consequently, faster behavioral responses. This is in-line with the suggestion that attention improves perception by controlling the phase coherence of ongoing neural oscillations before stimulus onset. Our results provide the first evidence indicative of a functional use of low frequency phase to improve neural representations, by attention.

## Acknowledgements

We would like to thank Laura Busse and Steffen Katzner for sharing the data they recorded from the two monkeys. We also thank Dirk Prüsse, Leonore Burchardt, and Ralf Brockhausen for technical assistance.

## Author Contributions

ST and ME designed the study. BZ and KM performed data analyses. ME, MRD, HAM, BZ and ST interpreted the data. BZ, ME, ST, HAM and MRD wrote the paper.

## Decleration of Interests

The authors declare no competing interests.

## Experimental Procedures

### Animal Welfare

The scientists in this study are aware and are committed to the great responsibility they have in ensuring the best possible science with the least possible harm to any animals used in scientific research (Roelfsema and Treue, 2014). All animal procedures of this study have been approved by the responsible regional government office (Niedersaechsisches Landesamt fuer Verbraucherschutz und Lebensmittelsicherheit (LAVES)) under the permit numbers 33.42502/08-07.02 and 33.9.42502-04-064/07. The animals were group-housed with other macaque monkeys in facilities of the German Primate Center in Goettingen, Germany in accordance with all applicable German and European regulations. The facility provides the animals with an enriched environment (incl. a multitude of toys and wooden structures (Berger et al., 2018; Calapai et al., 2017)), and natural as well as artificial light, exceeding the size requirements of the European regulations, including access to outdoor space. Surgeries were performed aseptically under gas anesthesia using standard techniques, including appropriate peri-surgical analgesia and monitoring to minimize potential suffering (Pfefferle et al., 2018).

The German Primate Center has several staff veterinarians that regularly monitor and examine the animals and consult on procedures. During the study the animals had unrestricted access to food and fluid, except on the days where data were collected or the animals were trained on the behavioral paradigm. On these days the animals were allowed unlimited access to fluid through their performance in the behavioral paradigm. Here the animals received fluid rewards for every correctly performed trial. Throughout the study the animals’ psychological and veterinary welfare was monitored by the veterinarians, the animal facility staff and the lab’s scientists, all specialized in working with non-human primates. The animals participating in this study were healthy at the conclusion of our study and were subsequently used in other studies.

### Behavioral Paradigm and Recording

Two monkeys were trained to perform a spatial attention task in which they had to detect a brief change in either the color or direction of one of two moving RDPs (Katzner et al., 2009). Each trial started when the monkey fixated its gaze on a central fixation point. After 150 ms, a cue appeared on the screen informing the monkey to which of the two upcoming stimuli it should attend to. After 500 ms, the cue disappeared and the RDPs were shown in the two visual hemifields. One of them was placed inside the receptive field (RF) of the neuron being recorded and the other was shown outside the RF, on the opposite side of the screen in the symmetric position relative to the fixation point. A brief color/direction change occurred after a random time between 500 and 3550 ms in the target or distractor stimulus. The monkeys were rewarded with a drop of juice if they successfully reported the target change and ignored the distracter change. In those trials in which no target change occurred, the monkeys had to continue holding the lever until the trial ended after 3550 ms following the onset of stimuli LFP and single unit signals were recorded from are MT of the two monkeys using a five-channel multi-electrode recording system (Mini-Matrix, Thomas Recording, and Plexon data acquisition system, Plexon Inc.). More details about the task are available in the original publication based on this data (Katzner et al., 2009).

### Preprocessing and Time-frequency Analysis

The recordings came from 27 sessions (7 from one animal and 20 from another). We chose a total of 31 recording sites for the time-frequency analysis from sessions containing at least 50 trials. This selection does not affect the main finding: incorporating the sessions with a small number of trials also led to a positive phase coherence modulation at the time/frequency with maximum phase coherence modulation (Figure S4). All analyses were carried out in MATLAB (Mathworks, Natick, MA). The LFP phases were aligned to correct the phase lags created by the recording hardware using the method suggested by (Nelson et al., 2008). LFPs reflect the summed neural activity across the synapses of a population of neighboring neurons (Buzsáki et al., 2012). Despite the retinotopic organization of MT, the receptive fields of neighboring neurons do not fully overlap, expanding the population receptive field compared to the individual neurons’ receptive fields. Therefore, even the spatial cue might evoke a response in the LFP despite being outside the neural RF. To avoid any contamination of the activity evoked by cue presentation, with the calculation of phase, our analysis window started from 100 ms after cue onset. We extracted the LFP signals coming from the 600 ms interval starting from 100 ms following the cue onset until 700 ms after it. To compute the PCM maps, LFPs were filtered into 4 Hz bins with steps of 1 Hz using the function eegfilt from EEGLAB toolbox (Delorme and Makeig, 2004). There were 5 frequency bins available for the time-frequency analysis, with the centers changing between 6-10 Hz and the 600 ms LFP from each trial was zero-padded by a 1000 zeros before the interval and 1899 zeros after it. Thus, the concatenated zeros had a length of more than three times of the analysis time window, to avoid any edge effect by the filter. To calculate the attentional index, spike density functions were computed by convolving a Gaussian kernel function (σ=30 ms) with the spike trains.

### Phase Coherence Measurement

For obtaining the phase coherence at a given time, the instantaneous phase of the LFP signal was extracted using Hilbert transform. Next, we projected the phases of trials into a unit-radius circle in order to achieve the circular distribution. Afterwards, we computed the phase coherence for the two subsets of attend-in and attend-out trials for each site. To calculate the attentional modulation of phase coherence for each site, we divided the difference between the attend-in and attend-out phase coherence by the summation of phase coherence for the two trial subsets. Next, we averaged these indices along sites. For the behavioral analysis, we used a similarity measure of each trial phase to the global mean phase at a specific time-frequency point for attend-in trials. Therefore, we chose a similarity measure which is derived for each trial by computing the magnitude of the circular vector summation of a trial’s phase with the global mean phase vector while both of them were normalized to unit. In this way, the maximum value of this similarity measure is 2 (when a trial’s phase is in the same direction as the global mean vector) and the minimum is 0 (when a trial’s phase is in the opposite direction of the global mean vector).

### Statistics

For testing the significance of attention’s effect on phase coherence, we performed a paired ttest across the phase coherence values coming from each site in the two attention conditions at every time-frequency pair. To correct for multiple comparisons, we controlled type I error with a false discovery rate (FDR) algorithm (Benjamini and Yekutieli, 2001). Hence, the time-frequency pairs with their p-values lower than the optimal p-value generated by the algorithm (here 0.016) were taken as significant. For the circular statistical analyses, we used the Matlab-based Circular Statistics Toolbox (Berens, 2009) and permutation test. For the permutation test, we shuffled the trials into two subsets of trials for 10,000 times and calculated the difference in phase coherence to generate a distribution. Then we assessed the location of the real phase coherence difference in this distribution to obtain p-values.

## Supplemental Information

**Figure S1.**
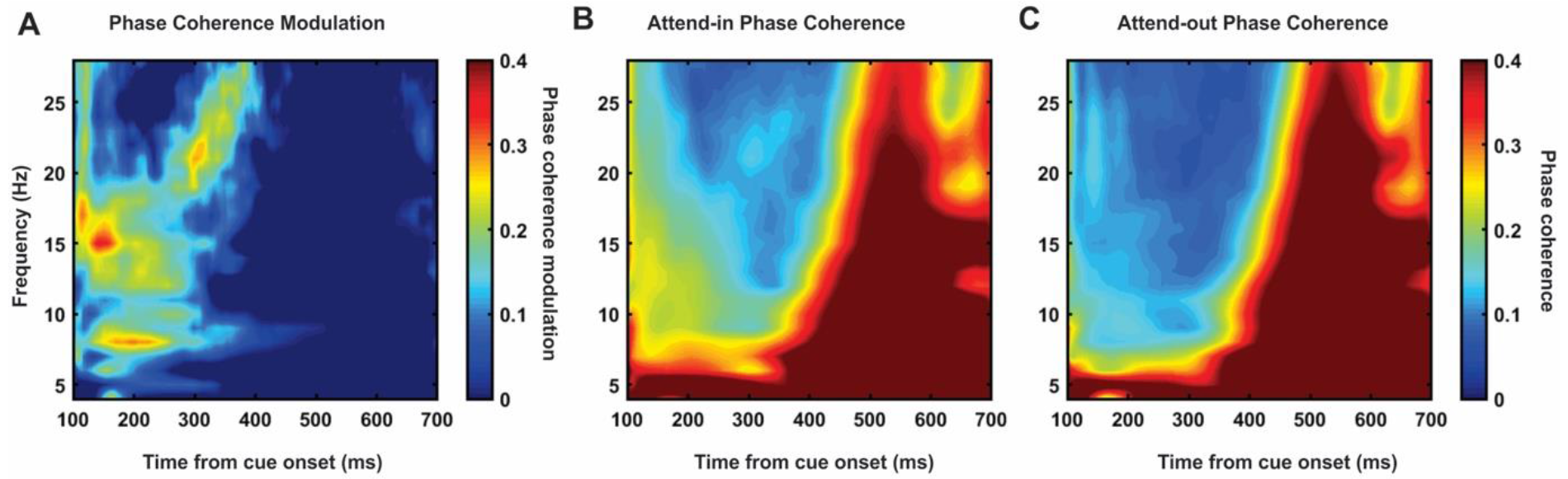
Full frequency range for phase coherence modulation. Related to Figure 1. (A) Phase coherence modulation for frequencies between 2 to 30 Hz. (B) Same as A, but for the average phase coherence across sites only for attend-in trials. (C) Same as B, but for the average phase coherence across sites only for attend-out trials.

**Figure S2.**
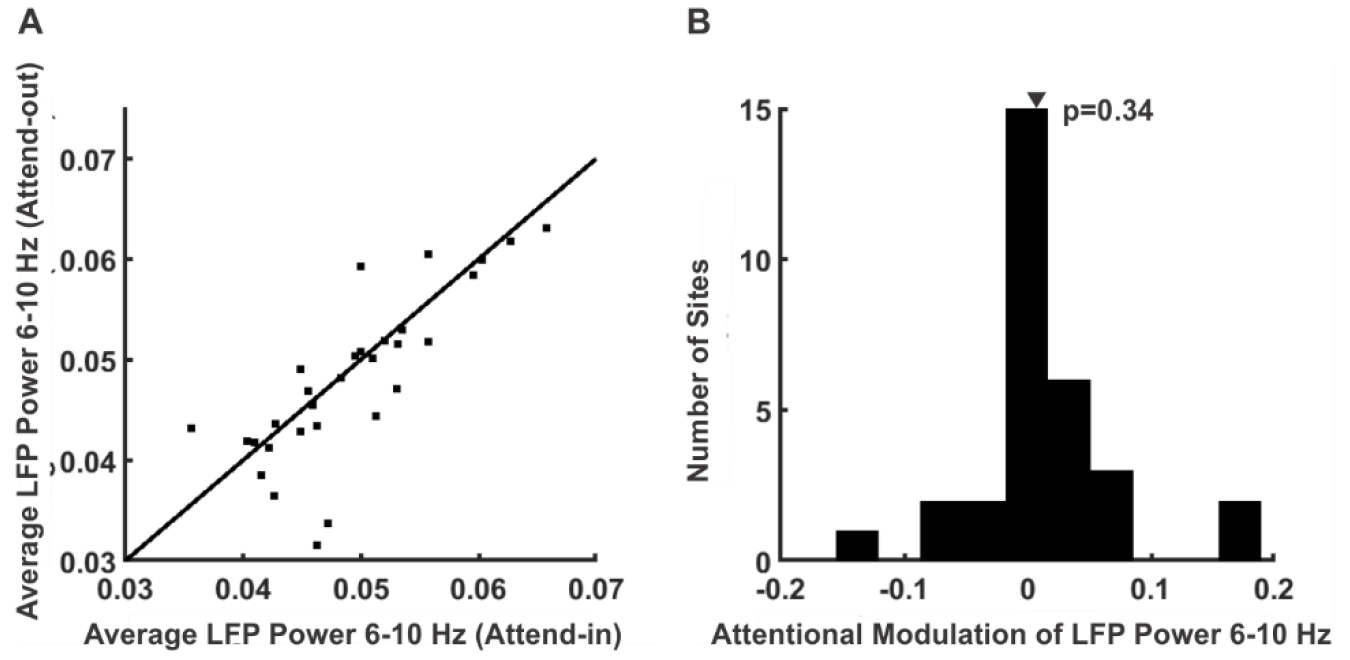
LFP power analysis within the pre-stimulus interval. Related to Figure 1. (A) The scatter plot shows the averaged LFP power across trials for 6-10 Hz band during 100-700 ms after cue onset. Each point corresponds to one site. The horizontal axis shows the average LFP power across trials for the attend-in subset of trials in each session and the vertical axis shows corresponds to the attend-out condition. (B) The histogram shows the attentional modulation of power in the same period showing no bias to positive values (p-value=0.34; ttest).

**Figure S3.**
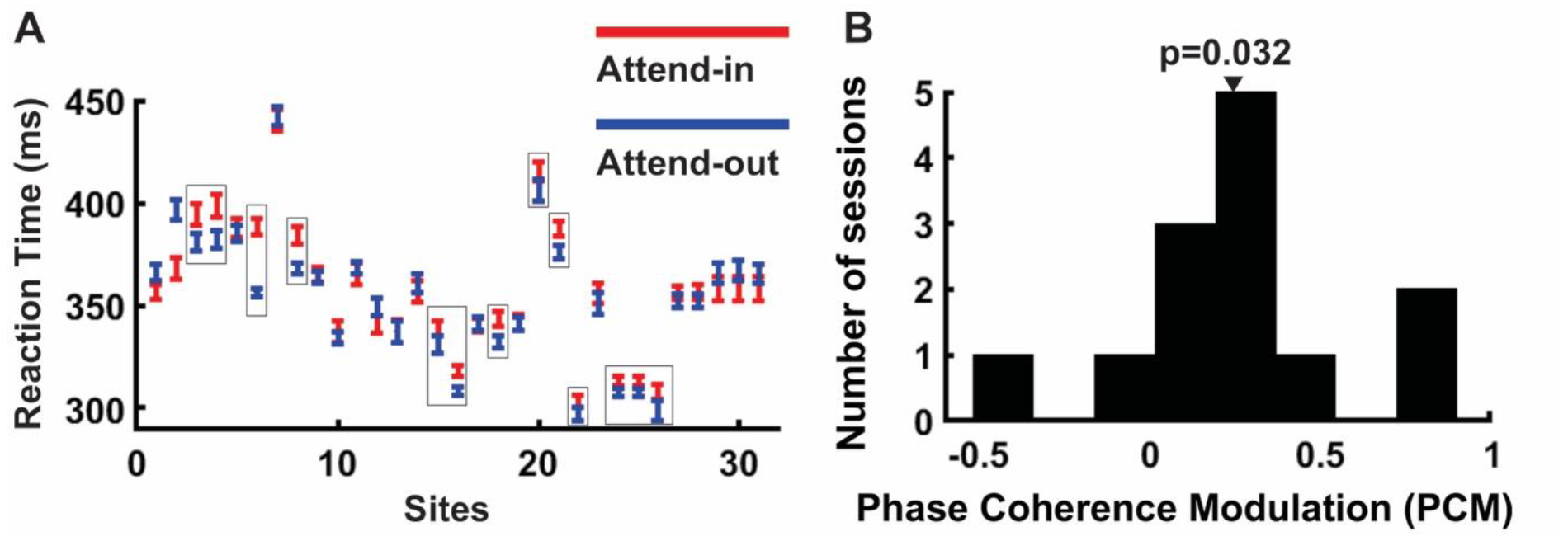
Control analysis addressing the difference of arousal level. Related to Figure 1. (A) X-axis indicates different sites and Y-axis represents the reaction time. Monkey responds faster in attend-out condition in sites indicated by rectangle. (B) Phase coherence modulation for sites indicated by rectangles in (A) (p-value<0.001; ttest). Bars show the standard error of the mean.

**Figure S4.**
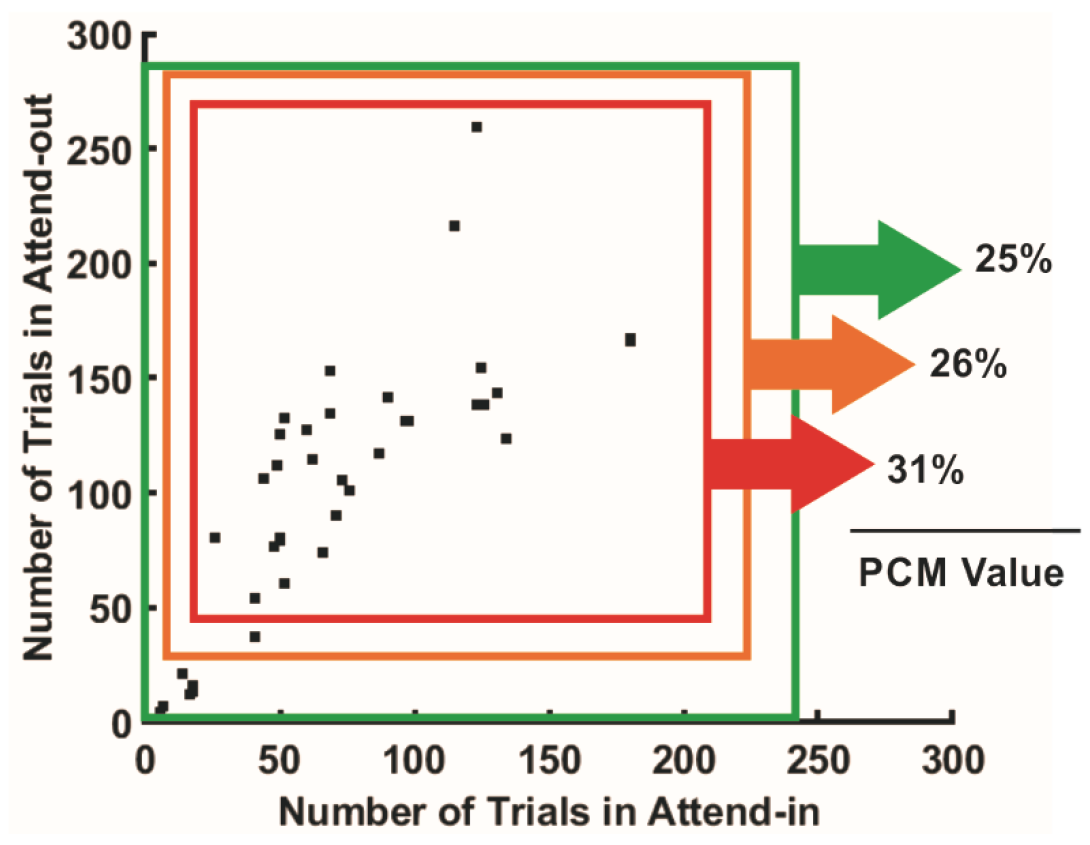
Phase coherence modulation at (200 ms, 8 Hz) including (excluding) the sites with low number of trials in both attention conditions. Related to Figure 1. Each point in the scatter plot corresponds to one site (from a total of 41 sites). The horizontal axis shows the number of trials in the attend-in condition and the vertical axis shows the attend-out condition. These results show that the PCM effect is observed independent of the selection of sites. The red square shows the set of sites focused for the analyses carried out here (31 sites).

